# GEPSi: A Python Library to Simulate GWAS Phenotype Data

**DOI:** 10.1101/2021.08.04.455085

**Authors:** Daniel A. Reidenbach, Avantika Lal, Lotfi Slim, Ohad Mosafi, Johnny Israeli

## Abstract

**Motivation:** Many computational methods aim to identify genetic variants associated with diseases and complex traits. Due to the absence of ground truth data, simulated genotype and phenotype data is needed to benchmark these methods. However, phenotypes are frequently simulated as an additive function of randomly selected variants, neglecting biological complexity such as non-random occurrence of causal SNPs, epistatic effects, heritability and dominance. Including such features would improve benchmarking studies and accelerate the development of methods for genetic analysis.

**Results:** Here, we describe GEPSi (**G**WAS **E**pistatic **P**henotype **Si**mulator), a user-friendly python package to simulate phenotype data based on user-supplied genotype data for a population. GEPSi incorporates diverse biological parameters such as heritability, dominance, population stratification and epistatic interactions between SNPs. We demonstrate the use of this package to compare machine learning methods for GWAS analysis.

**Availability and Implementation:** GEPSi is freely available under an Apache 2.0 license, and can be downloaded from https://github.com/clara-parabricks/GEPSi.

**Supplementary information:** Supplementary data are available online.

## 1 Introduction

Numerous computational methods have been developed to analyze Genome-Wide Association Study (GWAS) datasets, in order to identify germline genetic variants that are associated with phenotypic traits in a population. These include mixed linear models (Loh et al., 2015; Mbatchou et al., 2021), penalized logistic regression (Hoffman et al., 2013; Yang et al., 2020), decision trees (Botta et al., 2014; Schwarz et al., 2010), and neural networks (Arloth et al., 2020; van Hilten et al., 2020).

Due to the absence of ‘ground truth’ data where all the genetic variants associated with a phenotype are known with certainty, the small number of large-scale GWAS datasets available, and the protections surrounding access to such data, these methods are frequently developed using simulated genotype and phenotype data. This allows developers to benchmark their method on large and diverse datasets.

There are several existing tools to simulate genotypes based on diverse assumptions (Dimitromanolakis et al., 2019; Su et al., 2011). While phenotype simulation tools exist, most of these either randomly select causal SNPs (Meyer & Birney, 2018) or ask the user to specify them (Li & Li, 2008; Su et al., 2011). Further, it is common to assume a linear and independent additive genetic model, where the risk for an individual is a summation of risks for each causal SNP (Dimitromanolakis et al., 2019; Marigorta & Gibson, 2014; Meyer & Birney, 2018; Porter & O’Reilly, 2017).

However, causal SNPs for a trait are not randomly distributed throughout the genome, but instead are likely to directly or indirectly affect a common set of genes or pathways related to the phenotype. Moreover, various epistatic interactions between SNPs (Phillips, 2008) lead to non-linear relationships between genotype and phenotype. Although a few previous tools have included the option to specify non-additive effects for SNPs (Blumenthal et al., 2020; Tang Liu, 2019), to our knowledge there is currently no package dedicated to complex phenotypic simulations that incorporates nonlinear phenotype models, heritability, stratification and non-random selection of causal SNPs.

Here, we describe GEPSi, a python package to simulate phenotype data for a population based on given genotype data. GEPSi takes as input the real or simulated genotypes for all the individuals in a population. It simulates the phenotype of each individual based on user-specified parameters that control causal SNP selection, heritability, dominance, stratification and epistatic interactions between SNPs.

## 2 Workflow and Implementation

GEPSi requires two input files - a genotype matrix file in the ‘.raw’ format that specifies the genotype for each individual, and a SNP annotation file in the ‘.bim’ format that describes each SNP in the dataset. These files can be produced easily from other common formats using PLINK (Purcell et al., 2007).

GEPSi then simulates the phenotype for each individual causing the following steps (Figure 1A):

**Figure 1:**
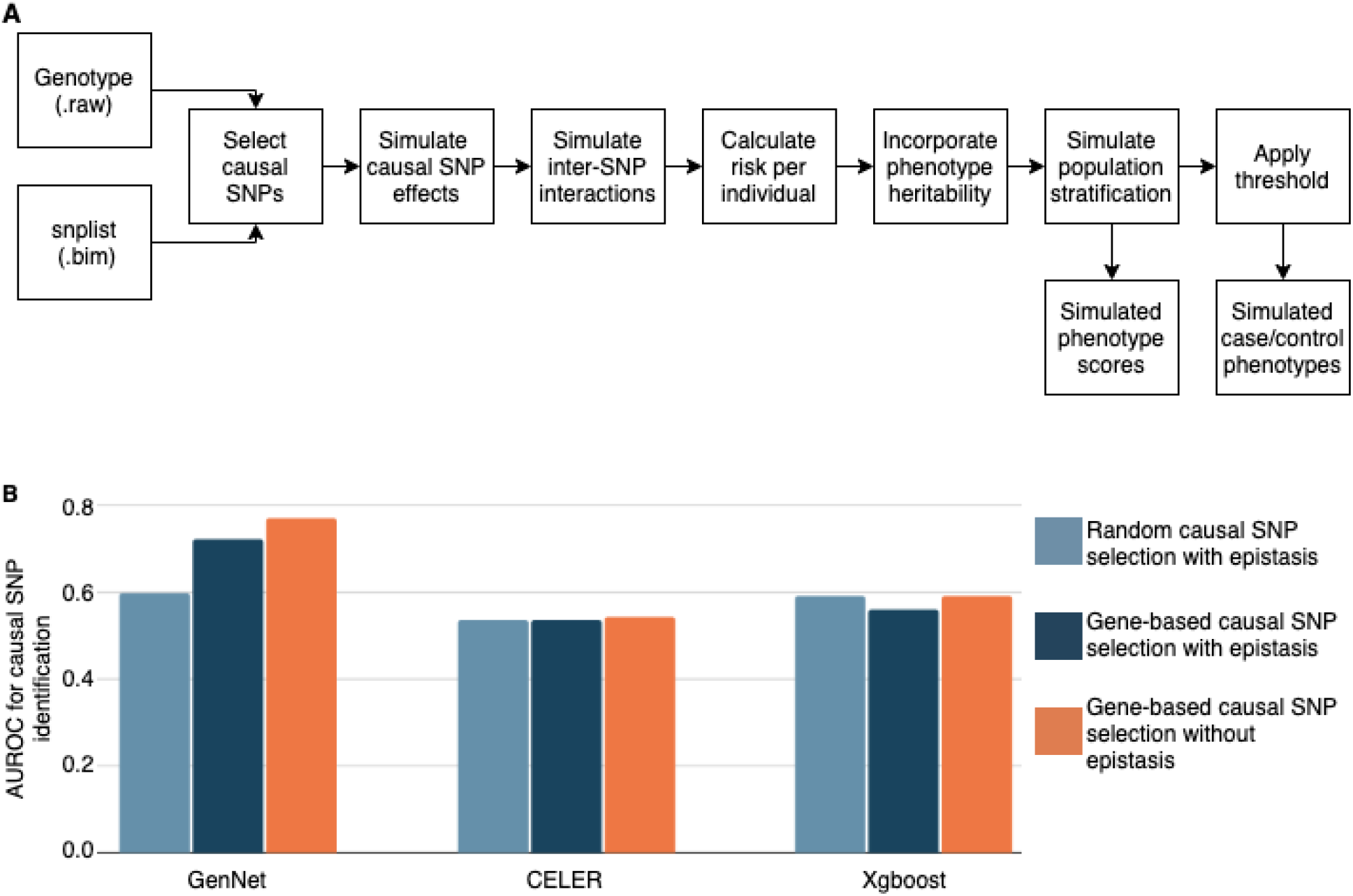
Benchmarking GWAS methods using GEPSi. A. Schematic of the GEPSi simulation procedure. B. Performance of three methods (GenNet, CELER and Xgboost) at identifying causal SNPs in three simulated GWAS datasets. While the genotype data was the same in all datasets, the phenotypes were simulated using GEPSi with three different settings.

1. A fraction of all available SNPs are selected to be causal. Selection can be random or gene-based. In gene-based selection, causal genes are first randomly selected and then causal SNPs are randomly selected from among the SNPs in the causal genes. This gene-based selection currently cannot be used to simulate causal intergenic variants.
2. A risk is assigned to each causal SNP by sampling from a normal distribution. Causal SNPs can be dominant, recessive, or dose-dependent with the user selecting the proportion belonging to each type.
3. Epistatic interactions are assigned to a fraction of the causal SNPs. Interactions between SNPs can be masking (in which one SNP masks or negates the effect of the other) or multiplicative (where one SNP increases or decreases the effect of the other).
4. The total genetic risk for each individual is calculated as the sum of the risks for all the causal SNPs based on their genotype.
5. The genetic risk for each individual is updated based on the heritability of the phenotype, according to the following equation:

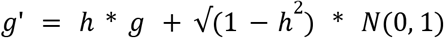

where *h* is the heritability and *g* the genetic risk score for the individual.
6. Population stratification may be incorporated by adding user-specified coefficients to the risk scores of individuals belonging to each group within the population.

User-controlled parameters include the causal SNP selection method, the number of causal SNPs of each type, the number of inter-SNP interactions of each type, the strength of inter-SNP interactions, the heritability of the phenotype, the stratification of the population into groups, and the proportion of cases and controls in the population The runtime of GEPSi with different settings is given in Supplementary Table 1.

## 3 Use Case

To demonstrate GEPSi, we applied HAPGEN2 (Su et al., 2011) to simulate chromosome 1 genotypes for a population of 10,000 individuals. We filtered 38,300 protein-coding SNPs for this analysis.

We then simulated phenotype data with three settings, each time selecting 1,290 of the 38,300 SNPs to be causal and setting 50% of the population to be cases while the other 50% were designated controls. The three simulations differed as follows:

1. Causal SNPs were randomly selected. Epistatic interactions were simulated for 20% of the causal SNPs.
2. Causal SNPs were clustered within 218 causal genes. Epistatic interactions were simulated for 20% of the causal SNPs.
3. Causal SNPs were clustered within 218 causal genes. Epistatic interactions were not included.

We then applied three machine learning methods - CELER Lasso (Massias et al., 2018), Xgboost, and GenNet (van Hilten et al., 2020) - to each simulated dataset, to classify individuals into cases and controls and to score the SNPs based on their association with the predicted phenotype (Supplementary Tables 2 and 3). All three methods were compared using the AUROC (Area Under the Receiver-Operator Characteristic) metric (Supplementary Tables 4 and 5).

GenNet, a deep learning method, performed best at identifying causal SNPs under all conditions (Figure 1B). While the performance of CELER and Xgboost was relatively constant across simulations, GenNet’s performance improved when causal SNPs were clustered within causal genes, rather than randomly selected (Figure 1B). Further, all methods performed slightly worse when epistatic interactions were included (Figure 1B). This example illustrates how GEPSi can be used to compare GWAS methods and to understand how genetic parameters influence model performance.

## 4 Conclusion

GEPSi enables easy benchmarking of GWAS software on simulated phenotypes based on a range of conditions and biological assumptions. It can also be used to benchmark other methods such as those to identify epistatic interactions (Niel et al., 2015).

In future, this framework can be extended to incorporate joint simulation of multiple phenotypes and epistatic interactions involving more than two SNPs or genes. Tissue-specific functional and pathway information, such as enhancer-gene maps (Nasser et al., 2021), can also be incorporated to simulate more realistic genotype-phenotype associations for the coding as well as non-coding genome.

