## Supplementary Material for "GEPSi: A Python Library to Simulate GWAS Phenotype Data"

### for:

Affiliations:
[1] Department of Electrical Engineering and Computer Sciences, UC Berkeley
[2] NVIDIA Corporation, Santa Clara, CA, USA
[3] NVIDIA Corporation, Reading, UK

**Supplementary Methods**

*Simulation of genotype data*
HAPGEN2 was used to simulate 10,000 genotypes based on phase 3 genotype data from the 1000 Genomes project. The simulated genotypes were filtered to 38,300 protein-coding SNPs on chromosome 1, by intersecting the SNP locations with exon locations from Gencode v19 (ftp://ftp.ebi.ac.uk/pub/databases/gencode/Gencode_human/release_19/gencode.v19.annotation.gtf.gz). All initial data was based on the GRCh37 human genome assembly.

*Training and testing models on simulated datasets*
Each simulated population was randomly split into training, validation and test sets of 8000, 1000 and 1000 individuals each. CELER, Xgboost and GenNet models were trained on the training sets, and the best model from each method was chosen based on validation set performance. All models were trained to predict the simulated phenotype of an individual (case vs. control) from the simulated genotype.

GenNet models were built with two layers. The first layer connects SNPs to genes, i.e. it takes as input the genotype of an individual at all SNPs, and learns weights to predict the activity of all genes. The second layer connects genes to output, i.e. it learns weights to predict the phenotype from the gene activities predicted by the first layer.

SNPs were scored based on their predicted association with disease by each method. CELER produces a coefficient for each SNP. For GenNet, each SNP was assigned a score by multiplying its weight in the first layer by the weight of the gene to which it was connected in the second layer. For Xgboost, the total cover score was used as the score for each SNP.

All three methods were compared on the test sets using the AUROC (Area Under the Receiver-Operator Characteristic) metric, which measures how well the scores assigned to SNPs by the model separate causal from non-causal SNPs.

**Supplementary Table 1: Runtime of GEPSi with different settings**

| SNPs | Population size | Genotype Raw Matrix Size | Memory setting | Genotype Runtime | Phenotype Runtime | Machine |
|---|---|---|---|---|---|---|
| 12,654 | 50K | 1.27 GB | Default | 3 min 27 sec | 12.1 sec | 16 Thread 32 GB RAM |
| 38,300 | 100K | 7.66 GB | Low Memory with Chunk size 20K | 51 min 51 sec | 1 min 30 sec | 16 Thread 32 GB RAM |

**Supplementary Table 2: Software versions used**

| Software | Software version |
|---|---|
| GenNet | 1.2.0 |
| Celer | 0.5.1 |
| XGBoost | 1.4.2 |

**Supplementary table 3: Software parameters used**

| Software | Parameters |
|---|---|
| GenNet | Adaboost, L1 = 1e-4<br>2 layers (SNP-gene and gene-phenotype) |
| Celer Lasso | L1 = 1e-3 for gene based simulation with epistasis, 1e-4 for others. |
| XGBoost | Max Depth = 8<br>Score = Total Cover |

**Supplementary Table 4: Simulation results - AUC for identification of causal SNPs**

| Simulation Type (38,300 SNPs, 10K Patients) | GenNet | CELER | Xgboost |
|---|---|---|---|
| Gene Based With Epistasis | 0.7224 | 0.5359 | 0.5590 |
| Gene Based Without Epistasis | 0.7690 | 0.5508 | 0.5911 |

| | | | |
|---|---|---|---|
| Random SNP selection with Epistasis | 0.5981 | 0.5500 | 0.5460 |

**Supplementary Table 5: Simulation results - AUC for identification of cases and controls**

| **Simulation Type (38,300 SNPs, 10K Patients)** | **GenNet** | **CELER** | **Xgboost** |
|---|---|---|---|
| Gene Based With Epistasis | 0.8449 | 0.8181 | 0.8790 |
| Gene Based Without Epistasis | 0.8425 | 0.7984 | 0.8515 |
| Random SNP selection with Epistasis | 0.8097 | 0.7923 | 0.7827 |
